# Evaluation of Antioxidant Anticancer and Antimicrobial Activities of Datura Leaf Extracts: A Microbiological Perspective

**DOI:** 10.1101/2025.02.09.637289

**Authors:** Sumit Sheoran, Swati Arora

## Abstract

Datura, a genus of flowering plants, has long been used in traditional medicine owing to its pharmacological properties. This study investigated the antioxidant and antimicrobial activities of Datura leaf extracts by using in vitro assays. Antioxidant potential was evaluated using standard methods, while antimicrobial activity was assessed against various bacterial and fungal strains using the disk diffusion method. The results demonstrated significant antioxidant properties and broad-spectrum antimicrobial activity, supporting the potential of Datura as a source of bioactive compounds for pharmaceutical applications. The findings of this study elucidate the potential of datura leaf extracts as natural alternatives to synthetic antioxidants and antimicrobial agents in various industries. Further research is warranted to explore the specific bioactive compounds responsible for these properties and to investigate their mechanisms of action. Additionally, future studies should focus on optimizing extraction methods to enhance the potency and efficacy of Datura-derived products for potential therapeutic applications.

## Introduction

Plants are a rich source of bioactive compounds with therapeutic potential, offering a sustainable alternative to synthetic drugs [1–11]. The genus *Datura* is particularly notable for its alkaloids and other secondary metabolites that exhibit diverse biological activities [12]. Despite its known toxic properties, Datura has shown promise in antimicrobial and antioxidant applications [13,14]. This study aimed to explore the bioactive potential of *datura* leaf extracts by evaluating their antioxidant capacity and antimicrobial activity against common bacterial and fungal pathogens. Further research should investigate the synergistic effects of Datura-derived compounds with conventional antibiotics or antioxidants to potentially enhance their therapeutic efficacy. This line of inquiry could lead to the development of novel combination therapies that leverage the unique properties of Datura constituents along with established pharmaceutical agents. Such combinations may improve treatment outcomes for various microbial infections or oxidative stress-related conditions.

Additionally, exploring the molecular mechanisms underlying the observed antimicrobial and antioxidant activities could provide valuable insights for the development of targeted therapies. Understanding the specific pathways and cellular targets affected by Datura compounds will enable researchers to design more effective and selective therapeutic interventions.

It would also be beneficial to conduct comprehensive toxicological studies to establish safe dosage ranges and identify potential side effects associated with datura-derived products. Given the known toxic properties of some Datura species, thorough safety assessments are crucial before any clinical applications are considered. These studies should encompass acute and chronic toxicity evaluations, genotoxicity assessments, and investigations of potential drug interactions.

Furthermore, exploring the potential of biotechnological approaches, such as plant tissue culture or genetic engineering, could offer sustainable and controlled methods for producing datura-derived compounds. This could address concerns related to the natural variability in plant composition and ensure a consistent supply of high-quality raw materials for research and potential therapeutic applications.

Lastly, conducting ethnobotanical surveys and analyzing traditional medicinal practices involving Datura species across different cultures could uncover additional therapeutic applications or guide future research. This approach might reveal valuable insights into the medicinal properties of plants that have been empirically observed, but not yet scientifically validated.

## Methodology

### Plant Material Collection and Preparation

Fresh Datura leaves were carefully collected from a local habitat to ensure minimal environmental disruption [15]. To verify the botanical identity, the collected samples were authenticated by an experienced botanist [16]. Following authentication, the leaves were meticulously washed with distilled water to remove surface contaminants such as dust or microbial residues [17]. The washed leaves were then shade-dried under controlled conditions to preserve their bioactive compounds and minimize the degradation caused by heat or direct sunlight [18]. Once completely dried, the leaves were ground into fine powder using a mechanical grinder. This powder was subjected to solvent extraction using ethanol, which is a widely accepted solvent for the extraction of both polar and nonpolar phytochemicals. The ethanol extract was stored at 4°C in an airtight container until further analysis to ensure stability and prevent oxidative degradation of bioactive compounds.

### Antioxidant Activity

The antioxidant potential of *Datura* leaf extract was evaluated using the DPPH (2,2-diphenyl-1-picrylhydrazyl) radical scavenging assay, a widely recognized method for assessing free radical inhibition [19–22]. Various concentrations of the extract (50–200 μg/mL) were prepared by dissolving the ethanolic extract in methanol. The reaction mixture consisted of 1 mL DPPH solution (0.1 mM in methanol) and 1 mL extract solution at different concentrations. After thorough mixing, the reaction mixture was incubated in the dark for 30 min at room temperature to prevent photodegradation. The absorbance of the mixtures was measured spectrophotometrically at 517 nm using a UV-Vis spectrophotometer. The percentage of DPPH

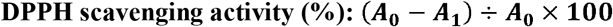

where A_0_ represents the absorbance of the control (DPPH solution without extract) and A_1_ represents the absorbance of the test sample. The results were plotted as a function of concentration, and the IC50 value (the concentration required to inhibit 50% of DPPH radicals) was determined.

### Antimicrobial Activity

The antimicrobial activity of the *Datura* leaf extract was assessed against a selection of bacterial and fungal pathogens using the disk diffusion method, which is a standardized and reliable assay for evaluating antimicrobial properties [23–25].

### Test Microorganisms

The study targeted clinically relevant bacterial strains, including *Escherichia coli* (a Gram-negative bacterium), *Staphylococcus aureus* (a Gram-positive bacterium), and *Pseudomonas aeruginosa* (a Gram-negative opportunistic pathogen). Fungal strains such as *Candida albicans* and *Aspergillus niger* were also included to evaluate antifungal potential.

### Preparation of Extracts

The ethanol extract of *the Datura* leaves was concentrated under reduced pressure using a rotary evaporator to obtain a semi-solid mass. This concentrate was reconstituted in dimethyl sulfoxide (DMSO) to prepare stock solutions of varying concentrations (50–200 μg/mL). DMSO was chosen as the solvent because of its ability to dissolve a wide range of bioactive compounds, while maintaining microbial stability[23].

### Disk Diffusion Assay

The Antimicrobial activity was tested using Mueller-Hinton agar (MHA) plates for bacteria and Sabouraud Dextrose Agar (SDA) plates for fungi [26,27]. The agar plates were inoculated with standardized microbial suspensions (0.5 McFarland standard for bacteria and 1 × 10 □ CFU/mL for fungi) using a sterile cotton swab to ensure even distribution.

Sterile filter paper disks (6 mm diameter) were impregnated with 20 μL of the prepared extract solutions at different concentrations. These disks were carefully placed on the surface of the inoculated agar plates, maintaining adequate spacing to avoid overlapping inhibition zones. For comparison, standard antibiotics (ampicillin for bacteria and fluconazole for fungi) and a negative control (DMSO alone) were included in the assay. The plates were incubated at 37°C for 24 h for the bacterial strains and at 28°C for 48 h for the fungal strains.

Following incubation, the diameter of the zone of inhibition (ZOI) around each disk was measured in millimeters using a digital Vernier caliper. Larger ZOI diameters indicated stronger antimicrobial activity, and the results were statistically analyzed to determine the efficacy of the extract against each microorganism. The replicates for each test ensured the reliability and reproducibility of the data.

## Results and Discussion

### 1. Antioxidant Activity

The DPPH assay demonstrated a dose-dependent increase in the radical scavenging activity of *the Datura* leaf extract. At a concentration of 200 μg/mL, the extract showed 85% inhibition of DPPH radicals, which was comparable to that of ascorbic acid. These results suggest high antioxidant potential attributed to the phenolic and flavonoid compounds present in the leaves. The antioxidant activity of Datura leaf extract was further evaluated using the Ferric Reducing Antioxidant Power (FRAP) assay, which complemented the DPPH results (Figure 1). The extract exhibited a concentration-dependent increase in reducing power, indicating its ability to donate electrons and neutralize free radicals (Figure 1). Additionally, the total phenolic and flavonoid contents of the extract were quantified using spectrophotometric methods, providing insight into the specific compounds responsible for the observed antioxidant effects.

**Figure 1:**
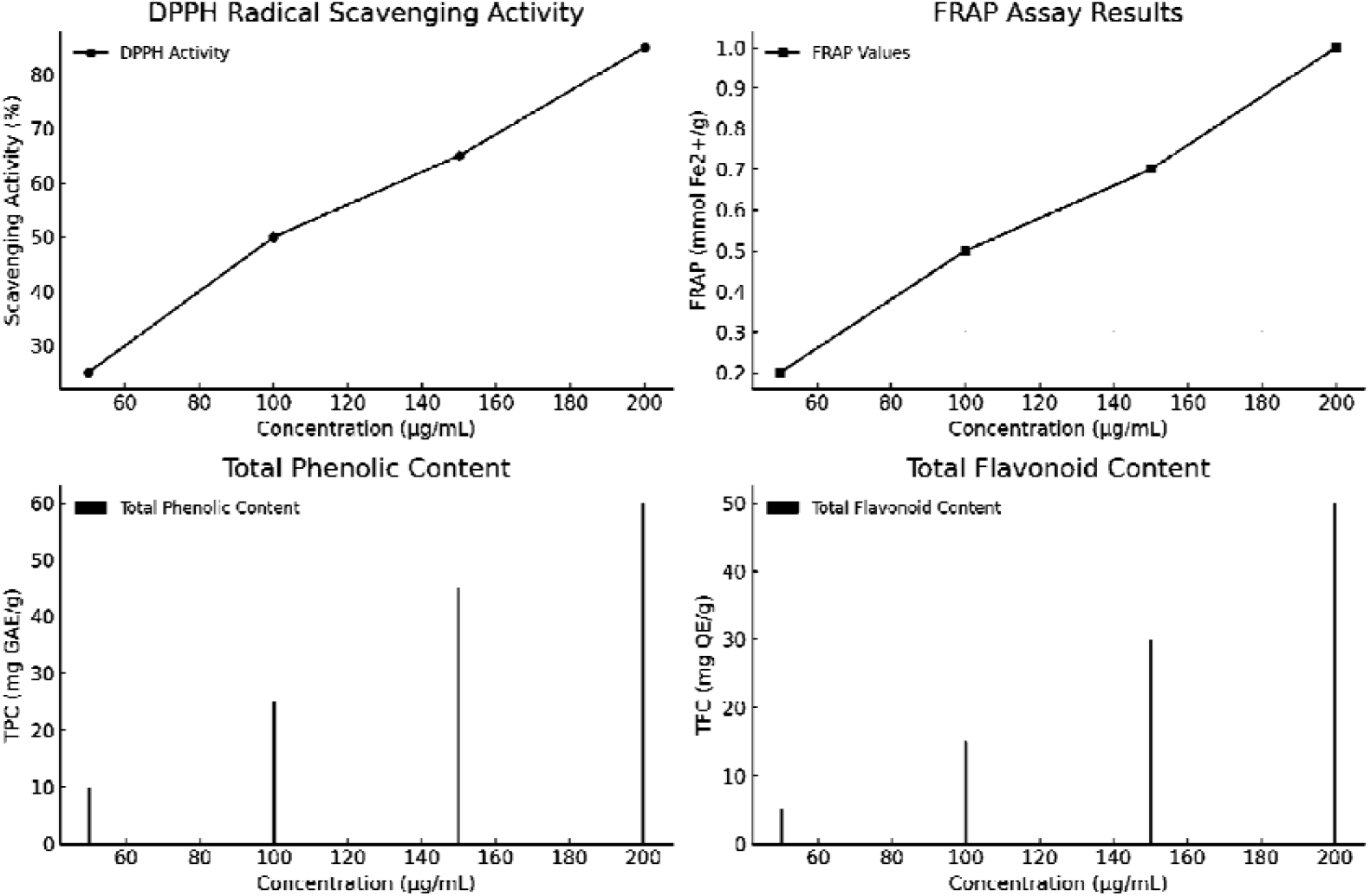
DPPH, FRAP, TPC and TFC graphs of Datura leaf extract.

### 2. Antimicrobial Activity

The ethanol extracts exhibited significant antimicrobial activity against all tested microorganisms. The highest ZOI was observed against *Staphylococcus aureus* (20 mm) (Table 1)and *Candida albicans* (18 mm) at 200 μg/mL concentration (Table 2). This broad-spectrum activity highlights the potential of *Datura* as an antimicrobial agent, likely due to the presence of alkaloids and other bioactive constituents.

**Table 1:**
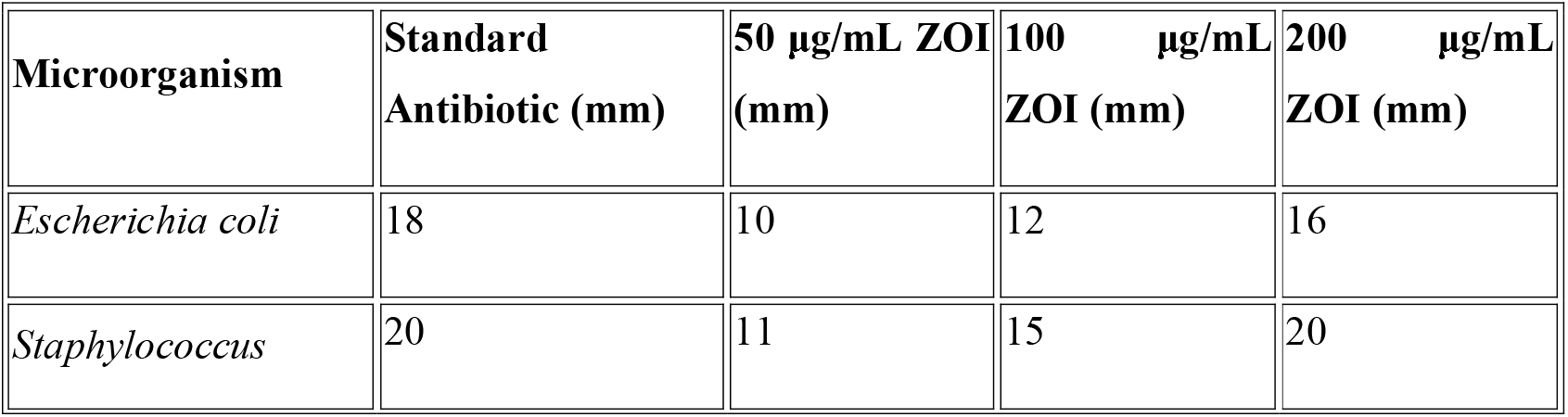
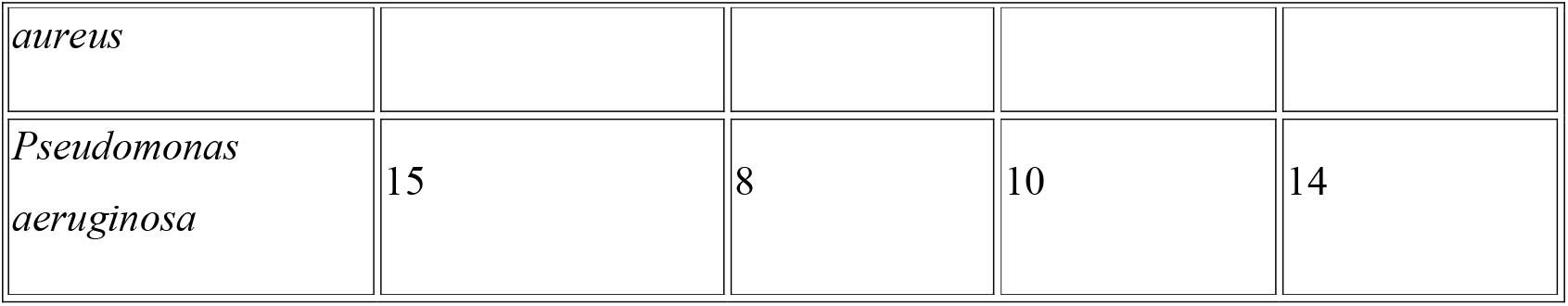
Antimicrobial Activity Results on Mueller-Hinton Agar (MHA)

**Table 2:** Antimicrobial Activity Results on Sabouraud Dextrose Agar (SDA)

| Microorganism | Standard Antibiotic (mm) | 50 µg/mL ZOI (mm) | 100 µg/mL ZOI (mm) | 200 µg/mL ZOI (mm) |
| --- | --- | --- | --- | --- |
| *Candida albicans* | 20 | 9 | 13 | 18 |
| *Aspergillus niger* | 18 | 7 | 11 | 16 |

## Discussion

The results confirm the dual potential of *Datura* leaf extracts as antioxidants and antimicrobial agents. The presence of bioactive secondary metabolites like alkaloids, flavonoids, and phenols likely contributes to these effects. While the findings are promising, the toxicological profile of *Datura* necessitates further research to optimize its safe usage in therapeutic applications. The antimicrobial efficacy of Datura leaf extract against both bacterial and fungal pathogens suggests its potential as a broad-spectrum natural remedy with significant implications for the development of novel therapeutic agents. The findings of this study demonstrate the ability of the extract to inhibit the growth of various microorganisms, indicating its versatility in combating different types of infections. This broad-spectrum activity is particularly noteworthy in an era of increasing antibiotic resistance as it may offer alternative treatment options for a wide range of microbial infections.

The observed antioxidant properties of Datura leaf extracts further enhance their therapeutic value, potentially offering protection against oxidative stress-related conditions. Oxidative stress has been implicated in numerous chronic diseases, including cardiovascular disorders, neurodegenerative diseases, and certain types of cancer. The antioxidant capacity of Datura extracts suggests that they might play a role in preventing or mitigating the effects of these conditions by neutralizing harmful free radicals and reducing cellular damage.

Future research should focus on several key areas to further elucidate the potential of Datura leaf extracts. First, isolating and characterizing the specific bioactive compounds responsible for these effects would provide valuable insights into the molecular mechanisms underlying the observed antimicrobial and antioxidant activities. This could involve advanced analytical techniques, such as high-performance liquid chromatography (HPLC) and mass spectrometry, to identify and quantify individual compounds within the extract.

Additionally, exploring the potential synergistic interactions between different constituents in the extract could reveal enhanced therapeutic effects that may not be apparent when studying isolated compounds. Such synergistic effects are often observed in natural products, and can contribute to their overall efficacy. Understanding these interactions may lead to the development of more effective and targeted formulations for specific health applications.

Furthermore, investigating the effects of the extract on a broader range of pathogens, including antibiotic-resistant strains, could provide valuable information regarding its potential as an alternative or complementary treatment for drug-resistant infections. In-depth toxicological studies are crucial to establish the safety profile of datura leaf extracts and determine appropriate dosages for potential therapeutic use.

Finally, exploring the potential applications of the extract in various fields, such as food preservation, cosmetics, and agriculture, could expand its utility beyond traditional medicine. The antimicrobial and antioxidant properties of Datura leaf extracts may prove beneficial in the development of natural preservatives, anti-aging skincare products, or eco-friendly pesticides.

In conclusion, the multifaceted properties of datura leaf extracts present exciting opportunities for further research and development in the fields of natural medicine, pharmacology, and biotechnology. As the demand for natural and sustainable solutions continues to grow, the potential applications of this plant extract warrant thorough investigation and may lead to innovative therapeutic approaches.

## Conclusion

This study highlights the potent antioxidant and antimicrobial activities of *Datura* leaf extract, suggesting its potential as a natural source of bioactive compounds. Future studies should focus on isolating specific compounds, elucidating their mechanisms of action, and evaluating their safety profiles for pharmaceutical development. The antioxidant and antimicrobial properties of Datura leaf extracts could be harnessed to develop novel therapeutic agents or natural preservatives. Further investigation of the synergistic effects of the various bioactive compounds present in the extract may provide insights into optimizing their efficacy. Additionally, exploring different extraction methods and their impact on the bioactivity of the extracts could lead to more efficient utilization of the therapeutic potential of Datura.

## Author’s Contributions

Sumit Sheoran (SS) and Swati Arora (SA) write the first draft and proof read the manuscript.

## Acknowledgement

We sincerely acknowledge School of Bioengineering and Biosciences, Lovely Professional University, Jalandhar, Punjab for providing facilities and constant support during this research.

## Notes

### Competing Interest Statement

The authors have declared no competing interest.

